# Emergence of prefrontal neuron maturation properties by training recurrent neural networks in cognitive tasks

**DOI:** 10.1101/2020.10.15.339663

**Authors:** Yichen Henry Liu, Junda Zhu, Christos Constantinidis, Xin Zhou

## Abstract

Working memory and response inhibition are functions that mature relatively late in life, after adolescence, paralleling the maturation of the prefrontal cortex. The link between behavioral and neural maturation is not obvious, however, making it challenging to understand how neural activity underlies the maturation of cognitive function. To gain insights into the nature of observed changes in prefrontal activity between adolescence and adulthood, we investigated the progressive changes in unit activity of Recurrent Neural Networks (RNNs) as they were trained to perform working memory and response inhibition tasks. These included increased delay period activity during working memory tasks, and increased activation in antisaccade tasks. These findings reveal universal properties underlying the neuronal computations behind cognitive tasks and explicate the nature of changes that occur as the result of developmental maturation.

## INTRODUCTION

Cognitive functions necessary for executive control such as working memory (the ability to maintain information in mind over a period of seconds) and response inhibition (the capacity to resist immediate responses and plan appropriate actions) mature relatively late in life, after adolescence^1–7^. This prolonged cognitive enhancement that persists after the onset of puberty parallels the maturation of the prefrontal cortex (PFC)^7–12^. Neurodevelopmental and psychiatric conditions such as ADHD, bipolar disorder, and schizophrenia are characterized by poor working memory and/or inhibitory control and they also manifest themselves in early adulthood^13–15^. Changing patterns of prefrontal activation between childhood and adulthood have been well documented in human imaging studies for tasks that require working memory^16–22^ and response inhibition^5,19,21,23,24^. Recordings from single neurons in these tasks have been recently obtained in non-human primate models, at different developmental stages^25–27^. Monkey maturation of working memory and response inhibition parallels that of humans^28^. The neurophysiological results are therefore highly informative about how mature cognitive functions are achieved in the adult brain. Many empirical changes observed in neurophysiological recordings between developmental states, however, remain difficult to interpret and it is unclear if systematic differences in neural activity at different developmental stages are causal to cognitive improvement, or incidental.

A potential means of understanding the nature of computations performed by neural circuits is to rely on Deep Learning methods^29,30^, which has been very fruitful in the analysis of the functions of neurons along the cortical visual pathways. Convolutional neural networks have had remarkable success in artificial vision and the properties of units in their hidden layers of have been found to mimic the properties of real neurons in the primate visual pathway^31–35^. It is possible to directly compare the activation profile of units in the hidden layers of artificial networks with neurons in the ventral visual pathway^36^. Deep learning models are thus being used to understand the development, organization, and computations of the sensory cortex^31–33^.

Another class of artificial networks models, Recurrent Neural Networks (RNNs) have been used recently to model performance of cognitive tasks and to study cortical areas involved in cognitive function^37,38^. The parameters of RNN architectures are fitted based on sequential data that can forecast future time series and are thus capable of exhibiting temporal dynamics resembling the time course of neural activity. RNNs can be trained to simulate performance of working memory and response inhibition tasks^39,40^, and have revealed, for example, how different neuron clusters represent different cognitive tasks^39^. We were motivated, therefore, to use RNNs at different stages of their training as a way of probing the changes that characterize maturation of real brain networks into a fully mature stage. Our analysis compared directly the levels of activity and temporal dynamics of single RNN units with those of single PFC neurons. We further sought to understand whether changes in activity we observed in PFC during brain maturation also emerged in the RNN network. Our work provides a framework for understanding the computations underlying cognitive maturation, simulating a wide range of cognitive tasks that may exhibit developmental differences, and generating predictions for further experimental study.

## RESULTS

Our approach compared the activity of real neurons recorded in the prefrontal cortex of adolescent and adult monkeys as they performed working memory and response inhibition tasks, and RNN units at different stages of training to perform the same tasks. Neural data were obtained from four macaque monkeys (*Macaca mulatta*) trained to perform two variants of the Oculomotor Delayed Response task (ODR and ODR plus distractor task) and three different variants of the antisaccade task differing in the timing of the cue onset relative to the fixation point offset (Fig. 1A). Data were obtained around the time of puberty (the “young” stage henceforth) and after the same animals had reached full maturity (the “adult” stage). In analogy, we examined responses of RNNs as the performance of the network improved during training in the exact same tasks.

**Figure 1.**
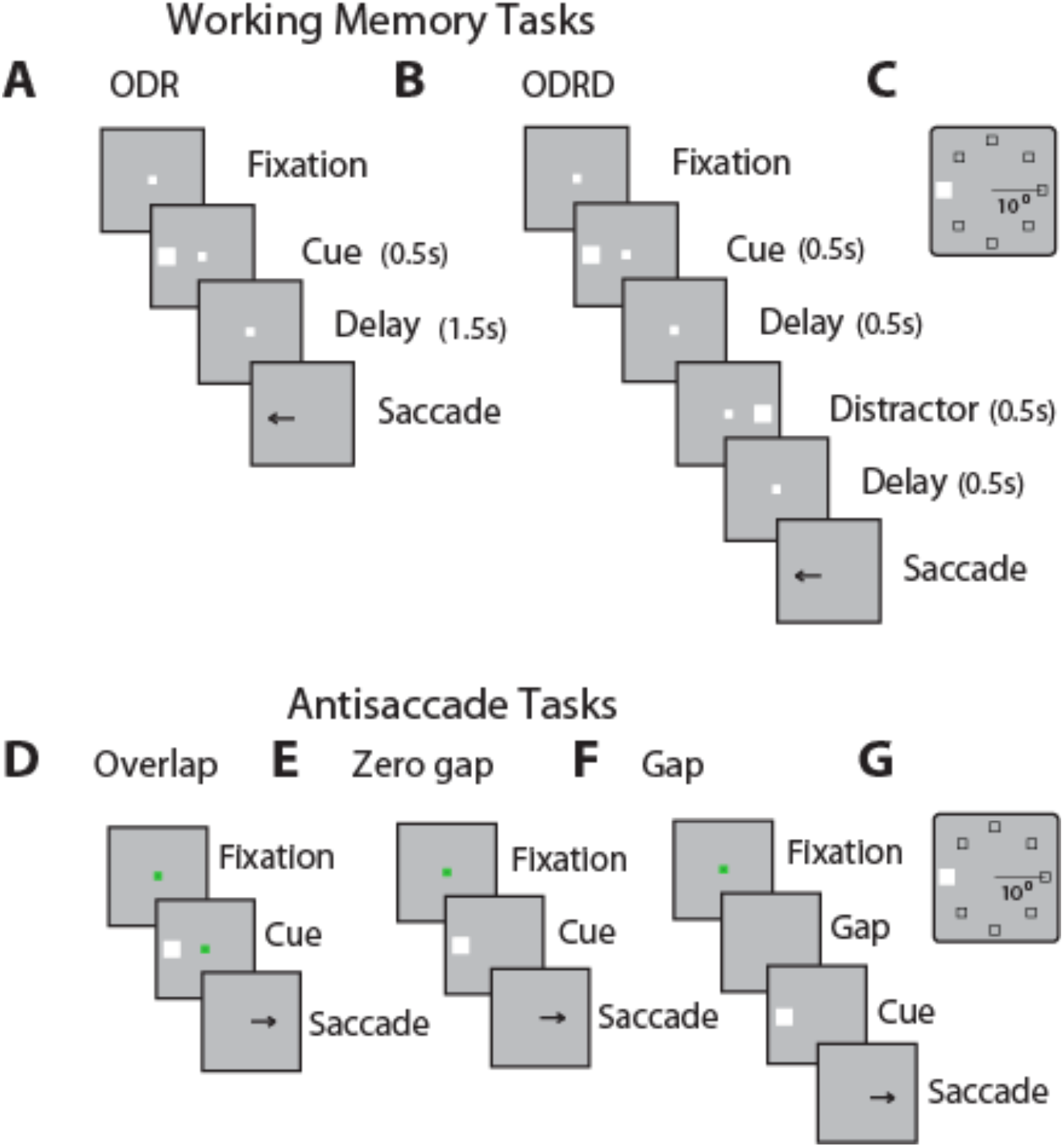
Behavioral Tasks. Successive frames illustrate the sequence of events in the working memory and antisaccade tasks simulated. A. Oculomotor Delayed Response (ODR) Task. B. ODR + distractor task. C. Possible locations of the cue in the ODR task. D. Overlap variant of the antisaccade task. E. Zero-gap variant of the antisaccade task. F. Gap variant of the antisaccade task. G. Schematic illustration of possible stimulus locations in the antisaccade tasks, which are the same as in the working memory tasks.

**Figure 2.**
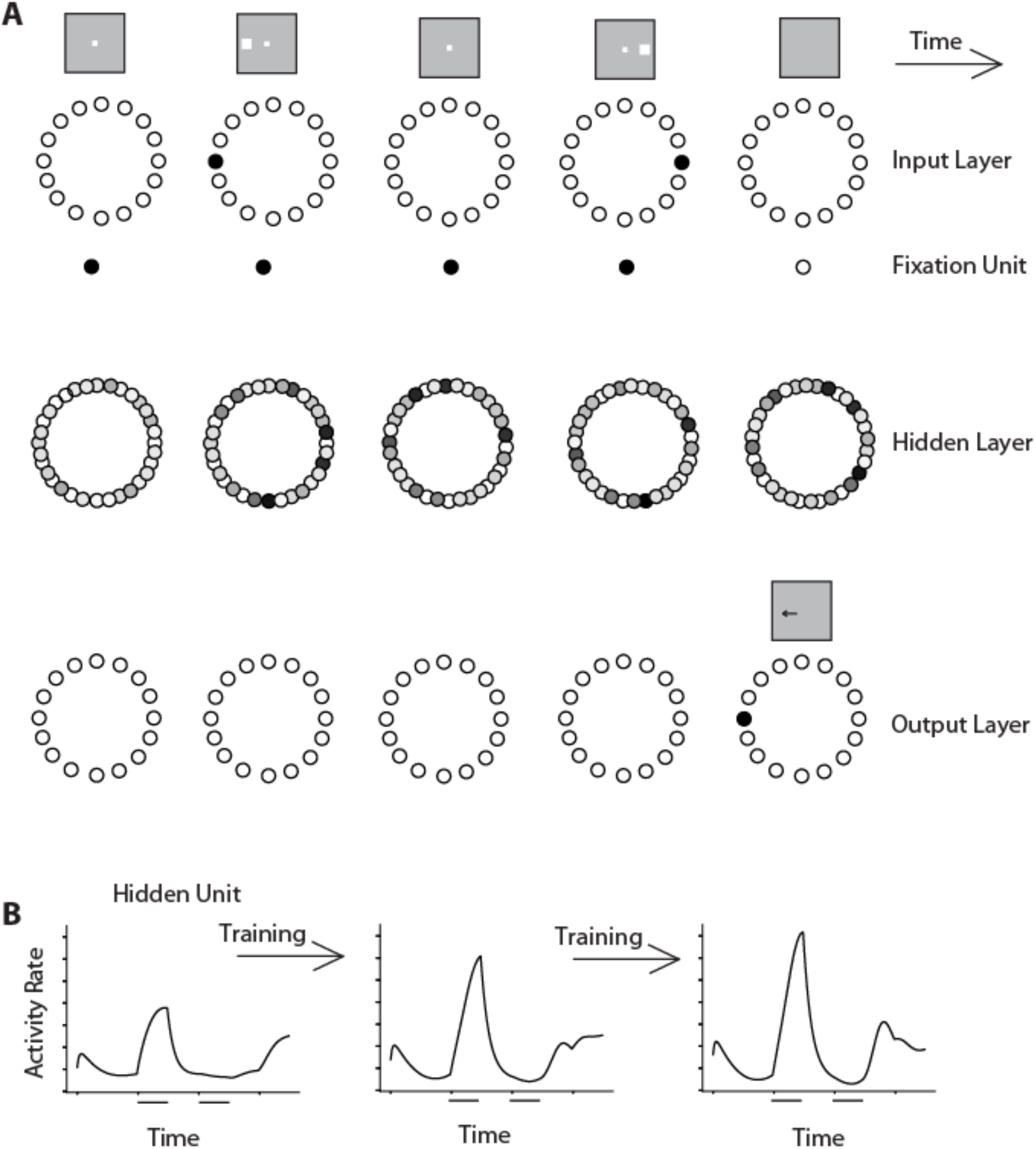
RNN network. **A.** Schematic architecture of input, hidden, and output layers of the network. Panels are arranged as to indicate successive events in time, in a single trial, across the horizontal axis. Appearance of the fixation point in the screen (top left panel) is simulated by virtue of activation of fixation units, a subset of the input units. Appearance of the visual stimulus to the left of the fixation activates input units in representing this location. Input units are connected to hidden layer units, and those to output layers units. In the ODRD task, the trained network generates a response to the by virtue of activation of the corresponding output unit. B. The firing rate of a single hidden unit is plotted as a function of time, during the duration of one trial. Successive panels now represent evolution of the units response as the training progresses.

We used an approach analogous to neurophysiological recordings to identify RNN units that respond to the task and study their responses to different task conditions. Such analysis of units in artificial neural networks has been called “artiphysiology” and offers direct insights on the role of neuronal activity in computation^36^. Use of RNNs allowed us to plot the entire time series of the unit’s activation during a trial in the task, and compare it with the Peri-stimulus Time Histogram (PSTH) of real prefrontal neurons. For each recurrent unit, we analyzed trials with the stimulus appearing at each of the eight locations, and calculated its firing rate during the cue and delay period. We identified individual units with responsiveness to the task, evidenced by a significant change in activity during stimulus presentation or delay period compared to the baseline period (t-test, p<0.05), and selectivity for different stimulus locations (1-way ANOVA test, p<0.05). The time course of their firing rate could then be plotted, in a manner similar to what we have done for real neurons (Fig. 3). Responses from responsive RNN units was averaged together to produce population activity. Importantly, we performed this analysis at different phases of the RNN network training, simulating different stages of the brain development. We identified three stages (early, middle, and mature), demarcated by transition points at which the network performed each task at a performance rate <35%, 35-65% and >65%, with the middle interval selected as to approximate the range of performance of adolescent monkeys in these tasks^25,27^.

**Figure 3.**
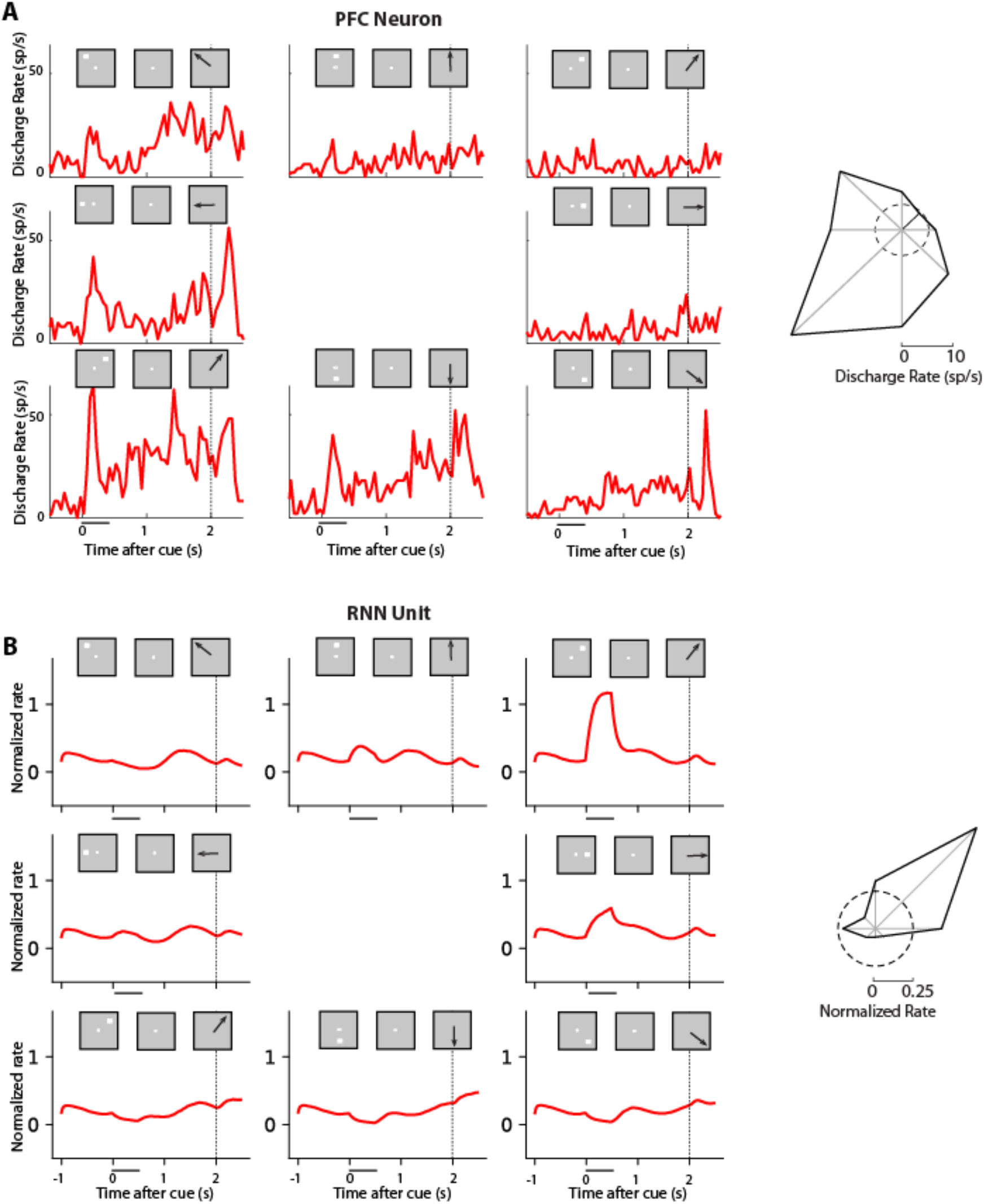
Activity of example units in working memory tasks. A. Firing rate of a single neuron in the monkey prefrontal cortex in the ODR task. Firing rate histograms are shown for eight locations, with plots arranged as to indicate the location of the stimulus in the screen. Insets above histograms indicate sequence of frames in the screen, which define the working memory task. Horizontal line indicates appearance of the cue; dotted vertical line the onset of the response. Polar plot depicts activity averaged over the entire cue period (from Zhou et al., 2013). B. Activity of unit from the adult stage of RNN for the exact same task presentations, as in A.

### RNN Activity in Working Memory Tasks

We first examined RNN activity in the simplest working memory task, the Oculomotor Delayed Response (ODR) task. As we have reported previously^41^, transition from the adolescent to the adult stage of maturation in monkeys is characterized primarily by an increase in activity of prefrontal neurons during the delay period of the ODR task, after the stimulus was no longer present (Fig. 4A). In contrast, responses to the visual stimulus itself remain essentially unchanged, at least relative to the baseline fixation period. We wished to test whether elevated activity during the delay period is a property of fully trained RNNs, particularly because the necessity of persistent neural activity in working memory itself has been a matter of debate in recent years^42,43^. Therefore, we identified RNN units with firing rate during the cue presentation period that was elevated relative to the fixation period (as was done for the analysis of prefrontal neurons) and compared their activity in the delay period across conditions and training stages. The transition of RNN unit activity from the early to the late stage of training resembled in many ways the maturation of the primate PFC. An increase in stimulus responses was evident early on, but this had essentially matured by the mid-trained stage (Fig. 4B). What continued to change until the fully trained stage was RNN unit activation that persisted in the delay period. The mean firing rate in the delay period differed significantly between phases (1-way ANOVA, F_2,331_=7.59, p=6.0×10^-4^ for the network units on which Fig. 4 was based). Average rate in each phase can be seen in Supplementary Fig. S1. The full time course of changes in activity can be seen in Supplementary Fig. S2.

**Figure 4.**
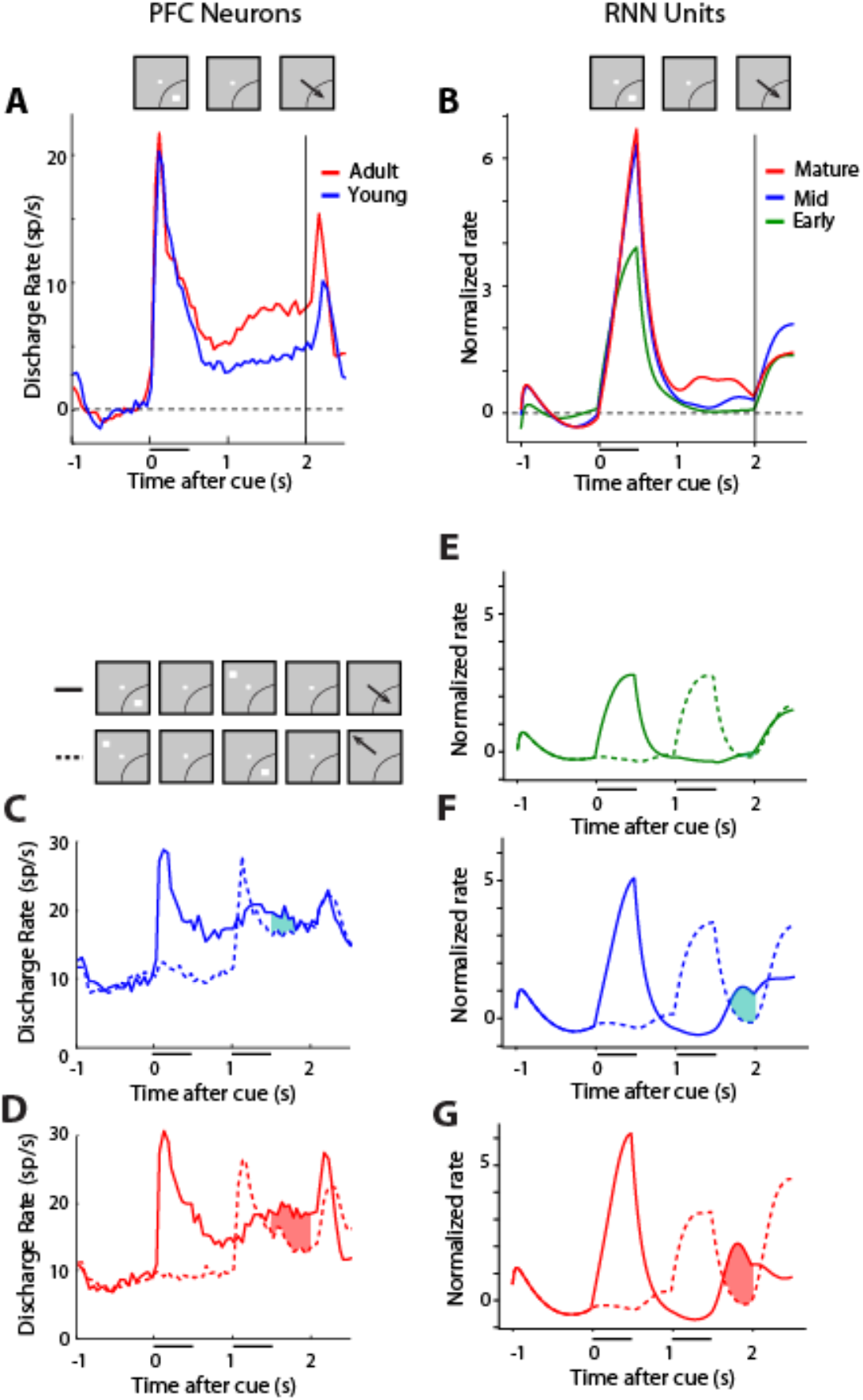
RNN and PFC activity in working memory tasks. A. Population discharge rate from young and adult PFC stages in the ODR task. Mean firing rate is plotted across neurons, after subtracting the respective baseline firing rate. Insets on top of the figure represent the sequence of events in the ODR task. Arc is meant to represent that the stimulus was presented at each neuron’s most responsive location. B. Mean rate of the RNN units responsive to the ODR task, during three developmental stages. C. Population discharge rate from the young PFC stage in the ODR+distractor task. Solid line represents appearance of a stimulus in the receptive field, followed by a distractor out of the receptive field. Insets represent sequence of events in the ODRD task. Dotted line represents appearance of a stimulus out of the receptive field, followed by a distractor in the receptive field. The difference between conditions indicated by the shaded line represents a measure of how well the distracting stimulus is filtered in the second delay period. D. Same as in C for adult data. E-G. Mean rate of RNN units plotted using the same conventions for three developmental stages. Neural data in panels A, C, D, from Zhou et al., 2016a.

Other RNN characteristics related to persistent activity also resembled neuronal responses. The number of selective RNN units for the different stimulus locations increased modestly but significantly between the different stages of training: 45% in the early phase, 51% in the middle, 56% in the mature phase (chi-square test, p=0.029). Across the population, RNN units trained to perform the ODR task also exhibited a population tuning profile that resembled that of the population of PFC neurons (Supplementary Fig. S3A-C). It was also possible to examine the activity of the exact same RNN units as training progressed. This analysis produced very similar results with the changes we saw in the averages (Supplementary Fig. S4). The activity of longitudinally tracked units paralleled the changes we saw across the population, for different training stages.

To gain insight in the changes of the temporal dynamics of unit activity as training progressed, we relied on a dimensionality reduction technique, termed dPCA, which has been used to understand dynamics of neuronal activity^44^, separating components relating to specific aspects of the task used. We processed the activity of RNN units as we did for neuronal activity and identified the top components capturing the temporal envelope of the response, selectivity for the visual stimulus location, and for mixture of time and stimulus (Fig. 5). The transition from the adolescent to the adult PFC was characterized by more robust separation of stimulus components during the delay period of the task, by themselves (Fig. 5A-B, middle panels) or in mixtures with time components (Fig. 5A-B, right panels). A very similar progression was observed in the RNN activity (Fig. 5C-E, middle panels), whereas early in training, stimulus components were separable only shortly after the cue appearance (Fig. 5C), in the mature network different stimuli were represented robustly through the end of the trial. However, some differences were also present; transient stimulus representation in the delay interval also emerged in the mature RNN (see Fig. 5E, right panel), in agreement with previous studies^45^, although this was absent in the PFC neural data. Cross temporal decoding similarly revealed that stability of the stimulus representation across the trial improved from the adolescent to the adult PFC, as did for RNN units (Supplementary Fig. S5).

**Figure 5.**
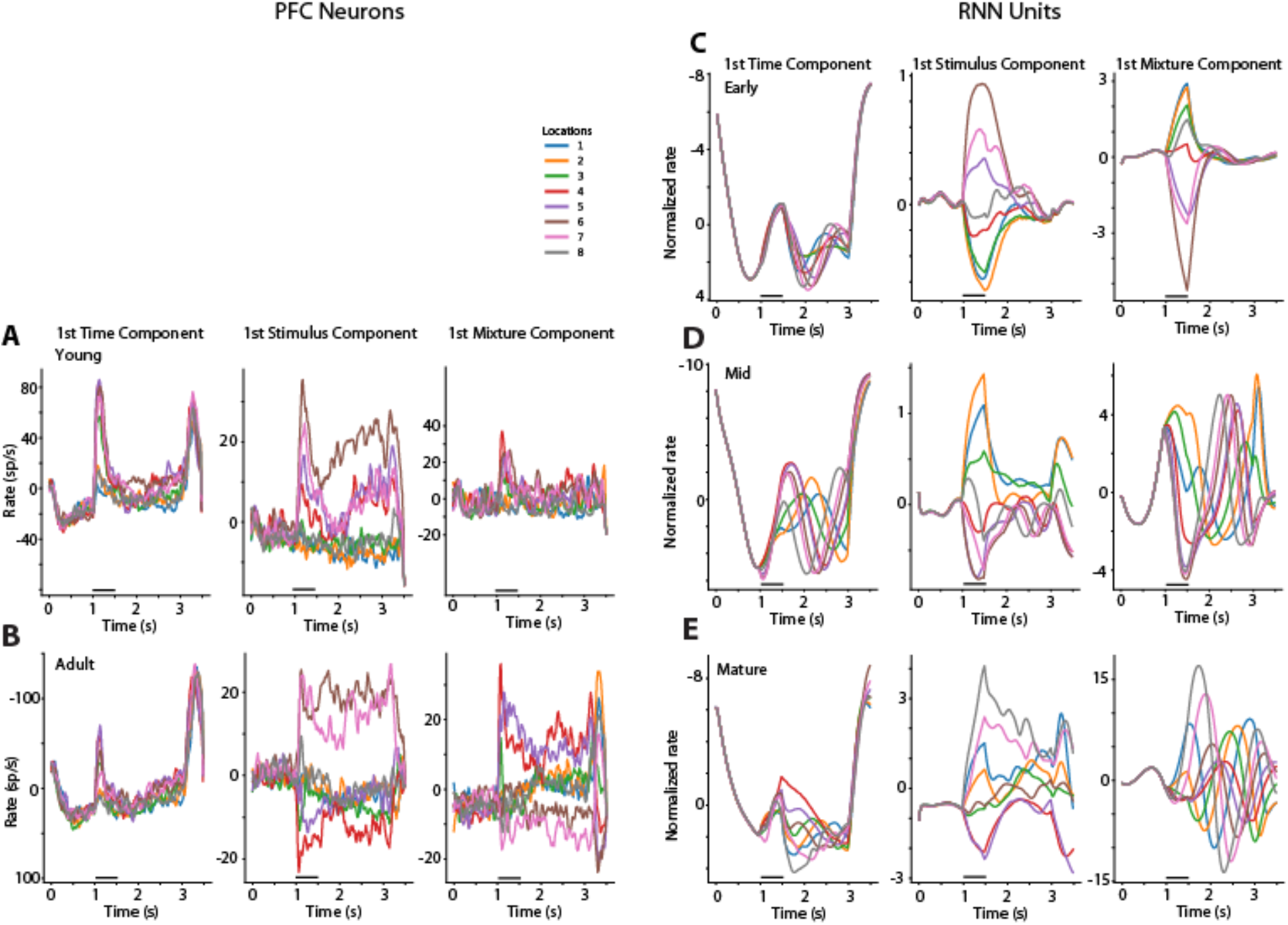
Demixed PCA Analysis. A. Data are plotted for PFC neurons recorded from the young monkeys. Panels from left to right represent the first time (condition-independent) component of dPCA analysis; the first stimulus-related dPCA component; and the first stimulus/time mixture dPCA component. B. Data from the Adult PFC. C-E, same analysis performed for RNN units in the early, mid-trained, and mature networks.

We also used a more challenging working memory task to probe the maturation of working memory ability, the ODRD task. In that task, after the original cue presentation that needed to be remembered, a distractor presentation followed that needed to be ignored over a second delay period (Fig. 1B). In neural activity, appearance of the cue in the receptive field generated a transient response, followed by persistent activity in the delay period of the task, which continued to be present, even after the distractor was presented (Fig. 4C-D). Appearance of the distractor in the receptive field also generated activation, however the relative difference between the activity generated by the cue and by the distractor in the second delay period of the task characterized the maturation of prefrontal neuronal activity (shaded area in Fig. 4C-D). This can be viewed as a measure of the prefrontal network ability to filter distracting activation^41^. In this case, too, we wished to test whether persistent activation of the RNN network was a result of training. In the early training stage, virtually no difference was present in the second delay period between cue and distractor-elicited activation (Fig. 4E). A difference between the two conditions began to emerge in the mid networks (Fig. 4F) and further grew in the fully trained ones (Fig. 4G). Tracking the activity of the same RNN units across training produced very similar results (Supplementary Fig. S2B-D). The mean firing rate across units in the second delay period differed significantly between phases (1-way ANOVA, F_2,579_=5.44, p=0.004; see also Supplementary Fig. S1B). In this case, too, the behavior of RNN units replicated the empirical pattern we have reported before in the PFC.

### RNN activity in response inhibition tasks

We proceeded to examine response inhibition, the ability to resist a response towards a prepotent stimulus, which is another cognitive domain that matures markedly between adolescence and adulthood. We relied on the antisaccade task, which involves presentation of a visual stimulus that subjects should resist looking at, but requires them to make an eye movement in the opposite direction (Fig. 1D-G). We have trained adolescent and adult monkeys in three different variants of the task of varying difficulty, by manipulating the relative timing of the cue presentation and offset of fixation point. The overlap variant was the easiest, as it allowed the monkeys to view the cue stimulus for 100 ms before both the fixation point turned off, which was the signal to the monkey to initiate the saccade (Fig. 1D). The gap variant (Fig. 1F) was the hardest as the fixation point turned off for 100 ms before the cue appeared (creating a “gap” of 100 ms when the screen was blank). In the absence of a fixation point where the subject can hold its gaze, it is much more difficult to resist making an erroneous saccade to the stimulus, than away from it. The zero gap variant involved turning off the fixation point simultaneously with the cue appearance and was intermediate in difficulty (Fig. 1D).

Neural responses in the antisaccade task mature considerably between the time of adolescence and adulthood^25^. In principle, improvement in performance in the antisaccade task may be achieved by a lower level of activation following the presentation of the visual stimulus, in essence filtering the representation of the prepotent stimulus towards which a saccade should be avoided or by increase of activity representing the target. The former alternative received some support, at least on the surface, by imaging studies which show decrease of prefrontal cortical activation between the time of adolescence and adulthood^46^. Somewhat unexpectedly, neurophysiological results supported the latter outcome. The most salient difference in neurophysiological recordings is an increase in activity in the adult stage for results synchronized on the onset of the saccade, even after subtracting the baseline firing rate. (Fig. 6A). We wished to test therefore if RNNs would capture this property. This was indeed the case (Fig. 6B). A progressive increase in the peak of activation was observed from early, to mid, to mature networks (Fig. 6B). This difference between stages was highly significant (1-way ANOVA for firing rate in 250ms after cue at early, mid, mature stage of gap task, F_2,638_=12.2, p=6.0×10^-6^).

**Figure 6.**
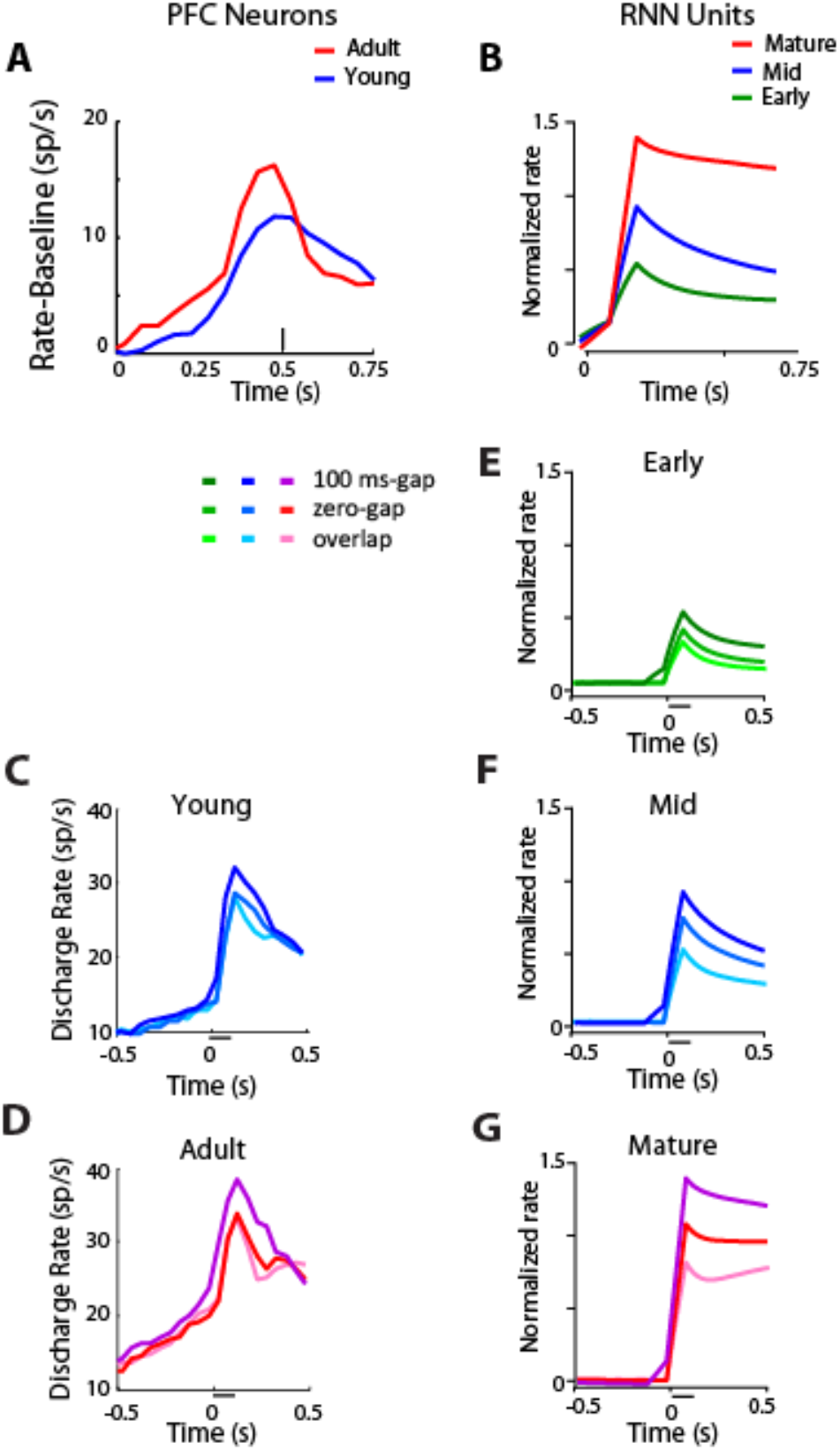
RNN and PFC activity in response inhibition tasks. A. Population discharge rate from young and adult PFC stages in the overlap variant of the antisaccade task, after subtracting the respective baseline firing rate. Data have been synchronized to the onset of the saccade, which is plotted here, at time 0.5 (indicated by vertical line). B. Mean rate of the RNN units responsive in the antisaccade task, during three developmental stages. C. Population discharge rate from the young PFC stage three variants of the antisaccade task: gap, zero-gap, and overlap. Horizontal line represents appearance of a stimulus in the receptive field. D. Same as in C for adult data. E-G. Mean rate of RNN units plotted using the same conventions for three developmental stages. Neural data in panels A, C, D, from Zhou et al., 2016b.

Interestingly, differences in firing rate elicited by different variants of the task at each training phase were also mirrored in the activity of RNN units. Higher firing rate was observed as a function of the task variant difficulty, with the gap variant eliciting the highest firing rate among prefrontal neurons. This was true for both the young stage (Fig. 6C) and the adult stage (Fig. 6D). The exact same pattern has been observed for RNN unit activation in the early (Fig. 6E), mid (Fig. 6F) and mature RNN networks (Fig. 6G). No neural theory to-date has been proposed to explain how these differences relate to task difficulty and performance. The current results demonstrate that emergent processes during the optimization process of artificial neural networks capture actual neural properties.

### Task Learning and Generalization

An important question for understanding the behavior of RNN networks is whether the network generalizes across tasks, i.e. represents similar task elements in the same manner, as the brain does, or creates entirely arbitrary associations. Previous work supports the idea that RNNs exhibit compositionality and perform similar task by activation of overlapping sets of units^39^. In the context of our tasks, we wished to understand whether units were tuned to the same stimulus location in the ODR and ODRD task. This was clearly the case, and this property persisted throughout the training regime (Supplementary Fig. S6).

A weaker correlation was observed between the best stimulus location that a unit exhibited in the ODR task and the Anti-saccade task (Supplementary Fig. S6B), also in agreement with neurophysiological data. This can be understood when one considers that the antisaccade task is fundamentally different than the ODR task than the ODRD is. It was notable that this divergence was amplified late in training, when the network mastered the task performance.

Analysis of the training phases revealed some “unnatural” behaviors as well. When all tasks were trained simultaneously, the RNN network mastered the antisaccade tasks much faster (Supplementary Fig. S7). The variants of the antisaccade task are conceptually very simple and require a straightforward mapping of weights in a naïve network: when a stimulus activates input unit *θ*, prepare a saccade at output position *θ*+*π*. However, this mapping is much more difficult to achieve in real brain networks, which are wired to perform saccades to the location of the visual stimulus.

### Network hyper-parameters

For the simulations we performed, we relied on parameters that were found to be effective for RNN networks learning to perform similar tasks^39,40^. However we wished to consider where the results were specific to this choice of hyper-parameters. We thus replicated these experiments by systematically varying the parameters of the recurrent neural networks (Supplementary Fig. S8). In networks that successfully acquired the tasks with different hyper-parameters, results were qualitatively similar and also resembled neuronal responses. We generally confirmed, however, that the hyper-parameters previously identified in the literature most closely resembled real neuronal responses for our results, too. In terms of weight initialization, randomized orthogonal weights produced best results, in contrast to diagonal initialization (Supplementary Fig. S8B), but random weights drawn from a Gaussian distribution also produced comparable results (Supplementary Fig. S8C). The number of the units of the network also affected network responses (Supplementary Fig. S8D-G), with results achieved by networks in the range of 128-512 units, most closely resembling neural results. Finally, the softplus activation function worked best compared to Relu, tanh, retanh, and power functions (Supplementary Fig. S8H-K), as it most closely resembles the nonlinearity function of real neurons.

## DISCUSSION

The use of artificial neural networks has exploded in recent years^47^. Convolutional neural networks have had remarkable success in artificial vision and the properties of hidden layers of artificial neural networks have been found to mimic the properties of real neurons in the primate visual pathway^31–35^. These results suggest that the human brain optimizes processing of visual information and that, after training, deep learning networks capture some of the same fundamental operations, allowing the use of such networks as scientific models^30^. The activity of individual units of these networks can be dissected, with techniques inspired from Neuroscience^36^. Neuroscience principles have also been instructive for the design of more efficient networks and learning algorithms^48^. In addition to convolutional networks, other architectures have had practical applications in Neuroscience questions, for example to uncover neuron spike dynamics, or encoding of elapsed time^49,50^. The activity of the prefrontal cortex has been investigated successfully with Recurrent Neural Network frameworks. Fully trained RNN models capture many properties of the prefrontal cortex, including its ability to maintain information in memory and to perform multiple cognitive tasks^39,40^.

Here, we capitalized on this approach to investigate a relatively unexplored aspect of prefrontal cortical function, its developmental maturation. In a series of recent studies, we have addressed for the first time the nature of changes that the prefrontal cortex undergoes between the time of puberty and adulthood^25–27,41^. These studies revealed a number of changes in activity between young and adult subjects, however the significance of changes at some task epochs relative to others and their relationship to behavior has been difficult to interpret. By using artificial RNNs, not explicitly modelled on any brain function, we show that the maturation of the primate prefrontal cortex optimizes processing necessary for cognitive tasks that require working memory and response inhibition.

### Parallel Neural and RNN Changes

The RNN model captured several activity properties from the mid-trained to the mature version which strongly paralleled the primate prefrontal development between the stage of adolescence to adulthood. In the ODR task, activity increased specifically during the delay period of the task, with little change in the cue presentation period itself. Stimulus representation was more separable during the delay period. In the ODRD task, the difference between delay period corresponding to the cue and distractor was also characteristic of development. The results also inform the debated of the basis of working memory and whether it depends on persistent discharges or not^42,43,51,52^. We saw that RNN units spontaneously develop working memory related activity, in agreement with previous studies^39,40^. Although it is possible for some RNN networks to perform working memory tasks without an overt increase in firing rate, more complex tasks do require persistent activity^40^. Our results support these conclusions and further indicate that the emergence of persistent activity is a developmental property that characterizes improved performance in mature networks.

Analysis of RNN activity also provided important insights about the neural basis of response inhibition. Prefrontal responses in the antisaccade task are characterized by differential levels of activity in task variants depending on difficulty, and parallel increases in maturation^25^. We now found that this pattern of responses emerges spontaneously in RNN networks, providing evidence again that the prefrontal activity optimizes the execution of the task. Improved performance in adulthood was not the result of more efficient filtering of the response to the stimulus, but rather stronger representation of the saccade goal.

### Model Limitations, Predictions, and Future Objectives

Although we have pointed out similarities in real and simulated networks, we do not wish to overstate this analogy. The RNN networks exhibited a number of “unnatural” behaviors, due to the unconstrained nature of their weights, and the complete lack of prior experience. For example, they were able to master the antisaccade task very easily, much faster than the working memory tasks. Although mapping an association between a stimulus and the opposite location is computationally straightforward, the brain is wired to direct saccades towards objects and not in diametric locations to them, which poses substantial challenges, particularly for the immature brain^28^. In essence, the real prefrontal cortex has already undergone a substantial shaping of its weights, which cannot always be captured by the RNN networks. In the ODR task, in addition to exhibiting persistent activity, they also maintained information in the delay period in a transient fashion, which is a documented property of RNNs trained to maintain information in short-term memory^45^. Both of these examples illustrate that RNNs are not expected to be a precise replica of the brain. The critical question is whether those patterns of activity that *do appear* in the RNNs also follow a similar trajectory as the maturing PFC. Our results document several aspects of activity that do in fact characterize improved performance in both the maturing brain and trained neural networks.

Use of RNNs to study neural processes opens up a wide array of opportunities for the study of cognitive development. In all simulations presented in the paper, we included an “early” training phase, meant to simulate the pre-adolescent state of the prefrontal cortex. By definition, the immature brain is unable to execute cognitive tasks at full capacity. For this reason teaching young subjects (humans and monkeys) tasks presents challenges. Scant neurophysiological findings are available from preadolescent subjects for this reason. Our findings make a number of predictions for immature PFC activity to be tested in future experiments: We posit that delay period activity is generated even at lower levels in working memory tasks, and it is less able to represent different stimuli; that distractors interrupt more effectively activity representing remembered stimuli; and that the antisaccade task elicits even lower levels of activity.

Further use of the RNNs training paradigm that we introduce here will also allow us to simulate a wide range of cognitive tasks and to identify ones that exhibit the widest contrasts between early and late training phases. These can then be implemented experimentally, and probed in terms of behavior and neurophysiology, allowing us to uncover hitherto unknown processes related to cognitive maturation.

## METHODS

### Behavioral and Neurophysiological Data

We relied on analysis of behavioral and neurophysiological results from monkeys performing working memory and response inhibition tasks, as these have been described in detail previously^25,41,53^. Briefly, the dataset was collected from four male rhesus monkeys (*Macaca mulatta*). All surgical and animal use procedures were reviewed and approved by the Wake Forest University Institutional Animal Care and Use Committee, in accordance with the U.S. Public Health Service Policy on humane care and use of laboratory animals and the National Research Council’s Guide for the care and use of laboratory animals. Development was tracked on a quarterly basis before, during, and after neurophysiological recordings allowing us to ascertain that data were collected during two stages: in adolescence, and adulthood^25,41^.

The monkeys were trained to perform two variants of the Oculomotor Delayed Response (ODR) task and three different variants of the antisaccade task (Fig. 1). The ODR task is a spatial working memory task requiring subjects to remember the location of a cue stimulus flashed on a screen for 0.5 s. The cue was a 1 ° white square stimulus that could appear at one of eight locations arranged on a circle of 10° eccentricity. After a 1.5 s delay period, the fixation point was extinguished and the monkey was trained to make an eye movement to the remembered location of the cue within 0.6 s. In the ODR with distractor variant (ODRD), a second stimulus appeared after a 0.5 s delay, followed by a second delay period. The monkey still needed to saccade to the original remembered stimulus. In the antisaccade task, each trial starts with the monkey fixating a central green point on the screen. After 1s fixation, the cue appears, consisting of a 1° white square stimulus that could appear at one of four locations arranged on a circle of 10° eccentricity for 0.1 s. The monkey is required to make a saccade at the location diametric to the cue. The saccade needed to terminate on a 5–6° radius window centered on the stimulus (within 3–4° from the edge of the stimulus), and the monkey was required to hold fixation within this window for 0.1 s. We used three different variants for the antisaccade task: overlap, zero gap, and gap, differing in the sequence of the cue onset relative to the fixation point offset (Fig. 1A). In the overlap condition, the cue appears first, and then fixation point and cue are simultaneously extinguished. In the zero gap condition, the fixation offset and the cue onset occur at the same time. In the gap condition, the fixation turns off and a 100 or 200 ms blank screen is inserted before the cue onset. Visual stimuli display, monitoring of eye position, and the synchronization of stimuli with neurophysiological data were performed with in-house software^54^ implemented on the MATLAB environment (Mathworks, Natick, MA). Behavioral and neural results were collected from the same animals in the young and adult stage.

Neural recordings were collected with epoxylite-coated Tungsten electrodes with a diameter of 250 μm and an impedance of 4 MΩ at 1 KHz (FHC Bowdoin, ME). Electrical signals recorded from the brain were amplified, band-pass filtered between 500 and 8 kHz, and stored through a modular data acquisition system at 25 μs resolution (APM system, FHC, Bowdoin, ME). Recordings analyzed here were obtained from areas 8a and 46 of the dorsolateral prefrontal cortex. Recorded spike waveforms were sorted into separate units using an automated cluster analysis method based on the KlustaKwik algorithm. Firing rate of units was then determined by averaging spikes in each task epoch. A total of 830 neurons from the adult stage, and 607 neurons from the young stage were used for subsequent analysis. In the ODR task, we identified neurons with significant elevation of firing rate in the 500 ms presentation of the cue, the 1500 ms delay period, and the 250 ms response epoch, after the offset of the fixation point. Firing rate in this period was compared to the 1 s baseline fixation period, prior to the presentation of the cue, and neurons with significant difference in firing rate were identified (paired t-test, p<0.05). Responsive neurons in the antisaccade task were identified based on significantly elevated responses in the 250 ms window following the onset of the cue compared to the fixation interval.

### Recurrent Neural Networks

We trained Leaky, Recurrent Neural Networks (RNNs) to perform multiple tasks simultaneously, following established procedures used to train RNNs to perform tasks used in neurophysiological experiments^39^. Implementation was based in Python 3.8, using the TensorFlow package. We used time-discretized RNNs with positive activity, modelled with the following equation (before descritization):

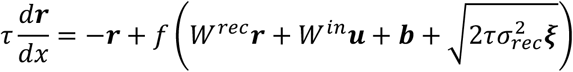

Where ⊤ is the neuronal time constant (set to 100 ms in our simulations), **u** the input to the network, **b** the background input, f the neuronal nonlinearity, **ξ** a vector of independent white noise process with zero mean and σ_rec_ the strength of noise (set to 0.05). Networks typically contained 256 units (results shown in main figures). Results from networks with numbers of units ranging from 64 to 1024 are shown in the supplementary material. We modeled the neuron nonlinearity based on the Softplus function

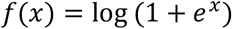

Results of networks with other functions are shown in the supplementary material, including ReLU (rectified, linear function),

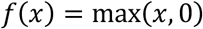

tanh (hyperbolic tangent),

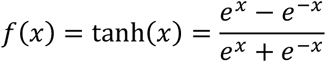

and retanh (rectified hyperbolic tangent):

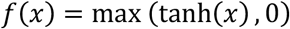

Output units, **z** read out the non-linearity from the network as:

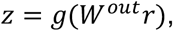

where *g*(*x*) is the logistic function

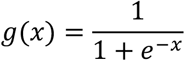

and *W^out^* the weights of units connected to the output units.

Our networks received three types of noisy input: fixation, visual stimulus location, and task rule. Weights were initialized with random orthogonal initialization (main figures), as well as diagonal, and random Gaussian values (supplementary material).

To train an RNN to perform the working memory tasks and anti-saccade tasks, we used a three dimensional tensor as the input to the network. This fully described the sequence of events in the six behavioral tasks used: ODR, ODRD, overlap, zero-gap, and two gap tasks (100 and 500 ms). The first dimension of the tensor encodes the noisy inputs of three types: fixation, stimulus location, and task rule. Fixation input was modeled as a binary input of either 1 (meaning the subject needs to fixate) or 0, otherwise. The stimulus is considered to appear at a ring of fixed eccentricity, and its location is fully determined by the angular dimension. Therefore, stimulus inputs consist of a ring of 8 units, with preferred directions uniformly spaced between 0 and 2π. The rule of the task was represented as a one-hot vector with a value of 1 representing the current task the subject is required to perform and 0 for all other possible tasks. The rule input unit corresponding to the current task was activated throughout the whole trial. Simulations involved 6 task inputs, therefore the first dimension of the tensor had a total of 1+8+6=15 inputs. The second dimension of the tensor encoded the batch size (number of trials). The third dimension encoded the time series for each trial.

A ring of 8 output units (plus one fixation output unit) similarly indicated the direction of gaze at each time point in the trial. While the fixation point was on, the fixation output unit should produce high activity. Once the fixation input was off, the subject had to make an eye movement (e.g. in the direction of the original stimulus in the ODR, or in the opposite direction in the antisaccade task), which was represented by activity in the network of tuned output units. The response direction of the network was read out using a population vector method. A trial is considered correct only if the network correctly maintained fixation (fixation output unit remained at a value >0.5) and the network responded within 36° of the target direction.

Each task can be separated into distinct epochs which duration was equal to the duration of the corresponding task in the neurophysiological experiments, described above. Fixation (fix) epoch is the period before any stimulus is shown, and lasted for 1 s. It was followed by the stimulus epoch 1 (stim1), which was equal to 0.5 s for the ODR and ODRD tasks, and 0.1 s fort the antisaccade tasks. If there are two stimuli separated in time, or if the stimulus and response are separated in time, then the period between the two is the delay epoch (1.5 s in the ODR task, 0.5 s in the ODRD task) and the second stimulus is epoch 2 (stim2, equal to 0.5 s). The period when the network should respond is the go epoch.

The RNNs are trained with supervised learning, based on variants of stochastic gradient descent, which modifies all connection weights (input, recurrent and output) to minimize a cost function L representing the difference between the network output and a desired (target) output^29^. We relied on the Adam optimization algorithm^55^ to update network weights iterative based in training data. For each step of training, the loss is computed using a small number M of randomly selected training examples, or minibatch. Trials representing all six tasks were included in a single minibatch during training of our networks. Trainable parameters, collectively denoted as ***θ*** are updated in the opposite direction of the gradient of the loss, with a magnitude proportional to the learning rate η:

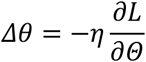

We found ability of the networks to master the task was quite sensitive to the value of η. This was set to 0.001 for most of the simulations included in the paper.

The activity of recurrent units was read out at discrete time points representing 20 ms bins. These can be compared with “Peri-stimulus Time histograms” of real neurons. Different stages of network training were used to simulate different developmental stages or task-training phases. For most analyses, we used both mixed training (the six tasks were randomly interleaved during training) and independent training (the network learned one task rule each time). We defined three “developmental” stages as the trial in which the network achieved <35%, 35-65% and >65% correct performance, respectively.

#### Demixed PCA and Decoding Analysis

Demixed Principal Component Analysis (dPCA) was performed, as we have described elsewhere^44^. The input is a four-dimensional array which represents number of neurons, discharge rate within a 20ms bin, stimulus locations, and trials for each stimulus condition. The method treats the responses of each neuron across time and stimulus conditions as one dimension, and then performs dimensionality reduction to determine components that correspond to stimulus and task variables.

Discharge rate across the length of the trial was calculated at every 200 ms window bins with an incremental step of 20 ms, and data from eight trials for the eight stimulus locations were then used for the analysis. Data from 256 RNN units were always used for RNN simulations. Decoding analysis was performed on the same dataset relying on an SVM classifier. The classifier was trained with a subset of trials at one window bin, and then tested with the remaining trials at the same window bin, and other window bins.

## Acknowledgements

Research reported in this paper was supported by the National Institute of Mental Health of the National Institutes of Health under award numbers R01 MH117996 and R01 MH116675. We wish to thank Balbir Singh and Jiajie Xiao for helpful comments on the manuscript.

## Code Availability

The code used to train, and analyze the units of the recurrent neural network is available at: https://github.com/maiziezhoulab/RNN_BrainMaturation

## Conflicts of Interest

The authors declare no competing interests

## Author Contributions

X.Z. and C.C. designed research; Y.H.L., and X.Z. performed simulations; J.Z. and X.Z. performed analysis of neural data. Y.H.L., X.Z. and C.C. wrote the paper.

## SUPPLEMENTARY MATERIAL

**Supplementary Figure 1.**
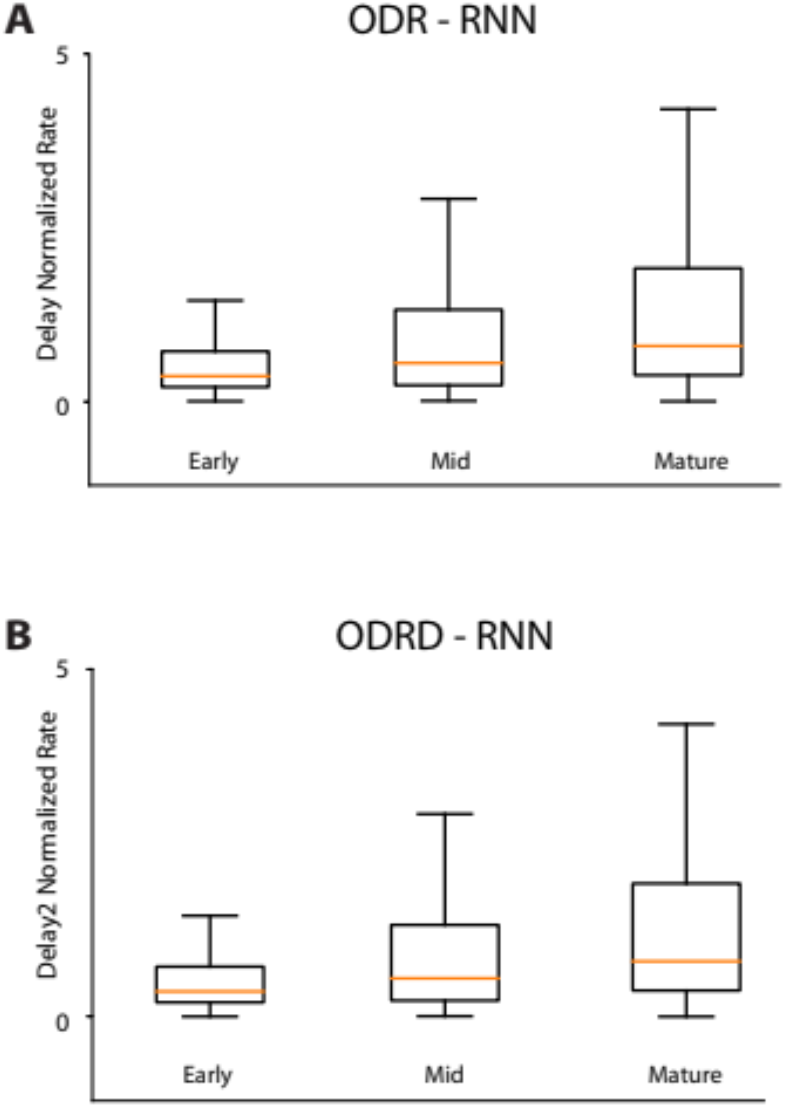
Delay period firing rate. A. Box plots represent median and interquartile range of firing rate of RNN unit activity during the delay period of the ODR task, computed separately for the early, mid, and late-training phase. B. Box plots of firing rate in the second delay period of the ODRD task.

**Supplementary Figure 2.**
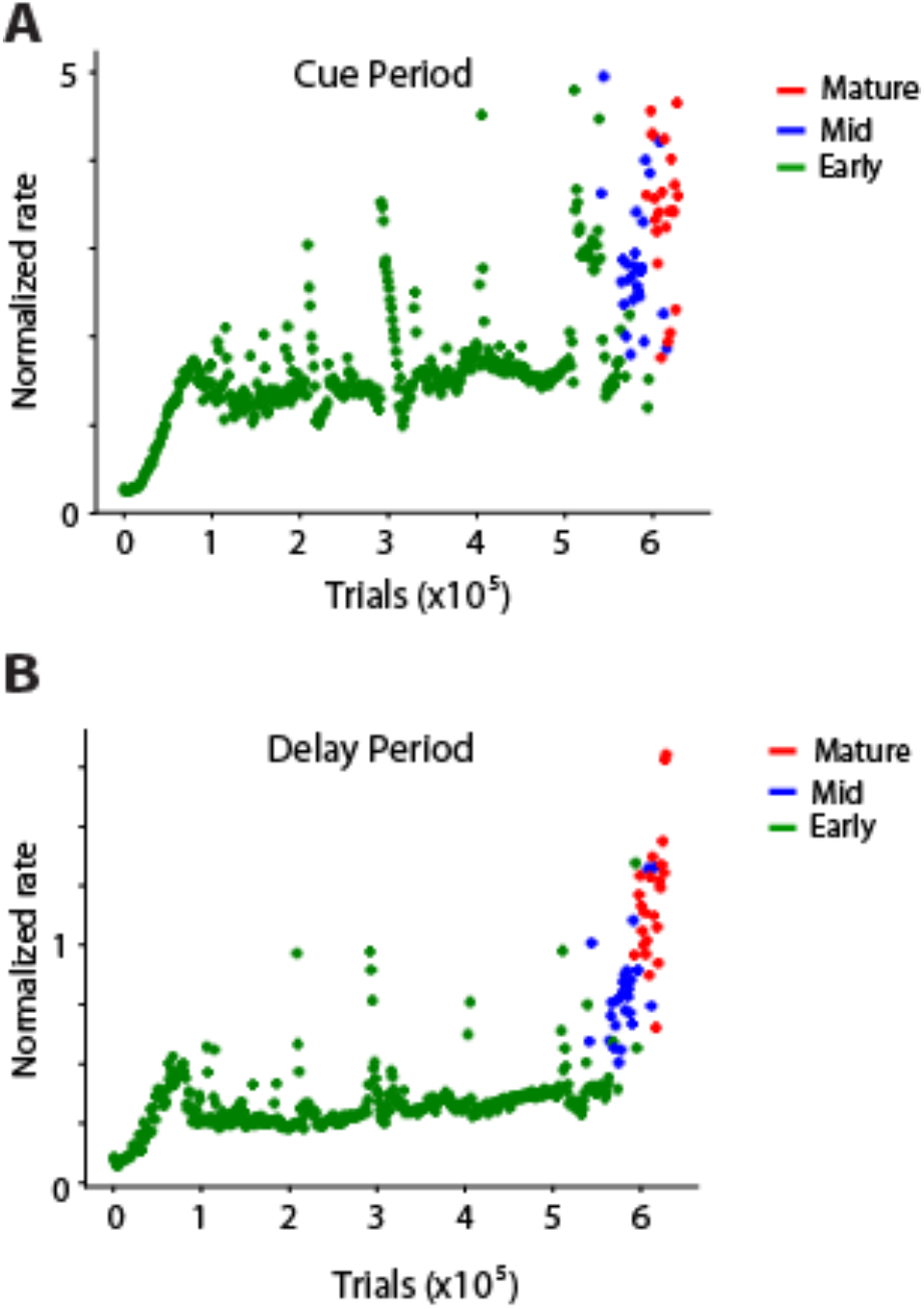
RNN activity during the course of training. A. Mean rate of the RNN units responsive to the ODR task during the cue period at successive trials. B. Mean rate of RNN units during the delay period.

**Supplementary Figure 3.**
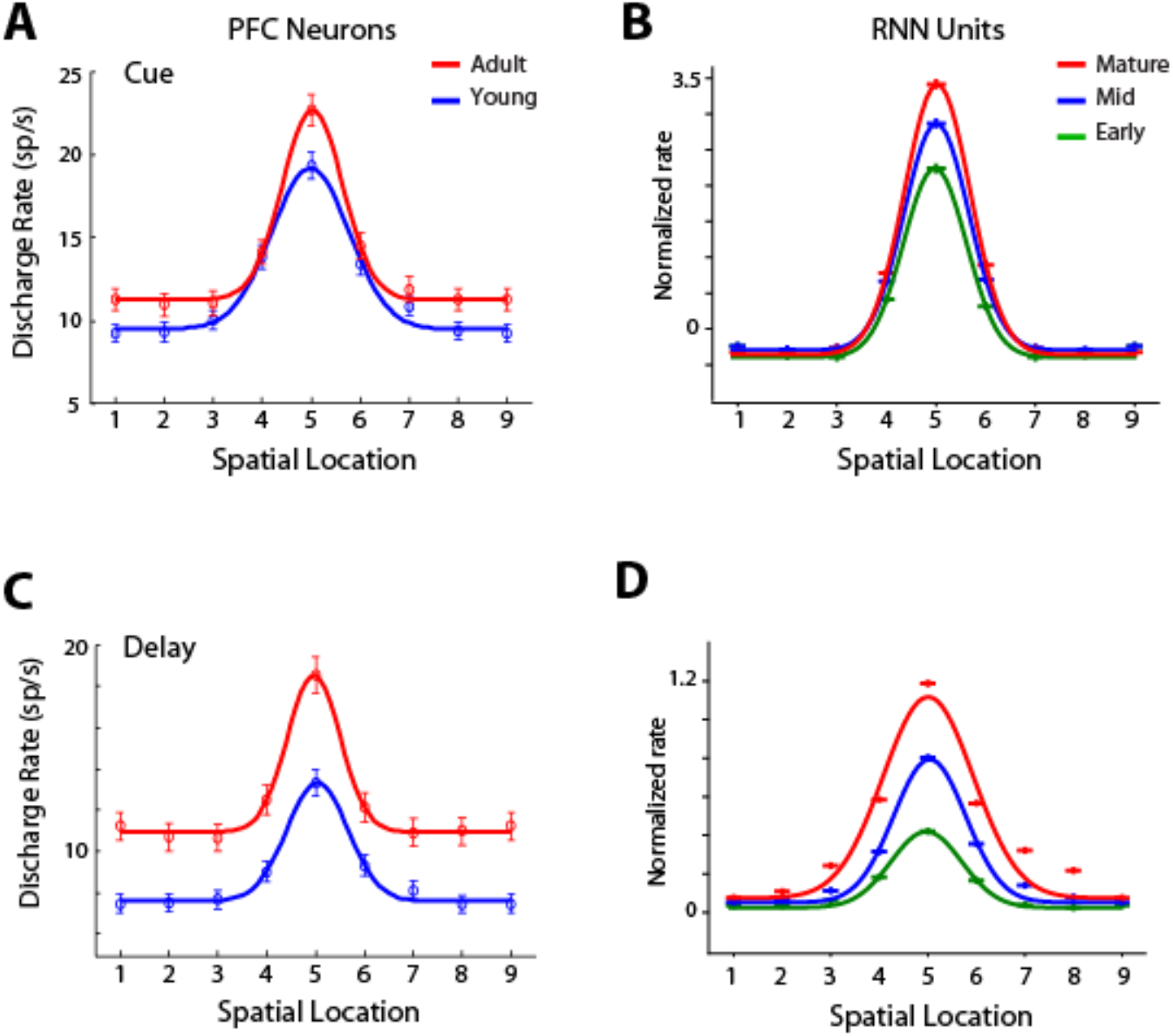
Tuning of RNN and PFC units. A. Population tuning curve from responsive PFC neurons in the young and adult stages of the ODR task constructed by rotating neuronal responses to the cue presentation at different spatial locations so that the best location of each neuron is represented in the graph’s center location (labeled 5). Curve represents best Gaussian fit. Error bars represent standard error of the mean across single neuron responses. B. Population tuning curve of RNN units responsive to the cue presentation in the ODR task, during three developmental stages. C. Population tuning curve from responsive PFC neurons during the delay period of the ODR task. D. Mean rate of RNN units plotted using the same conventions for three developmental stages. Neural data in panels A, C, from Zhou et al., 2016a.

**Supplementary Figure 4.**
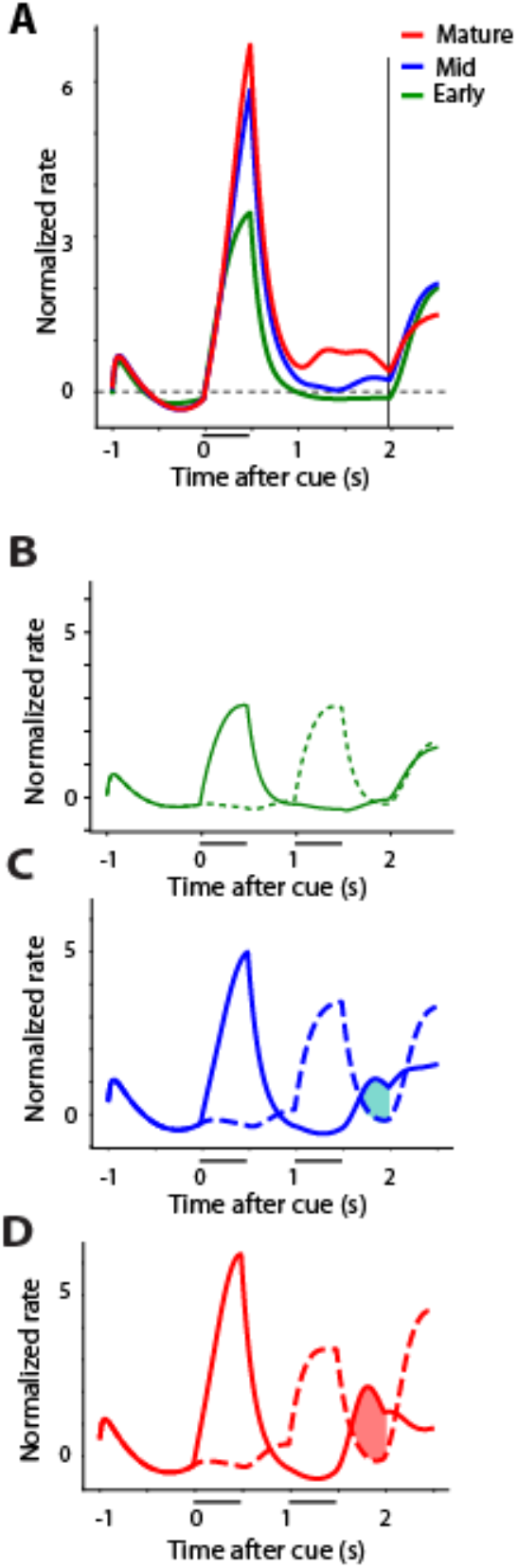
Activity of the same RNN units during training in working memory tasks. A. Firing rate of RNN units responsive to the ODR task, during three developmental stages. B-D. Mean rate of a the same RNNs units across developmental stages plotted using the same conventions as in Figure 4.

**Supplementary Figure 5.**
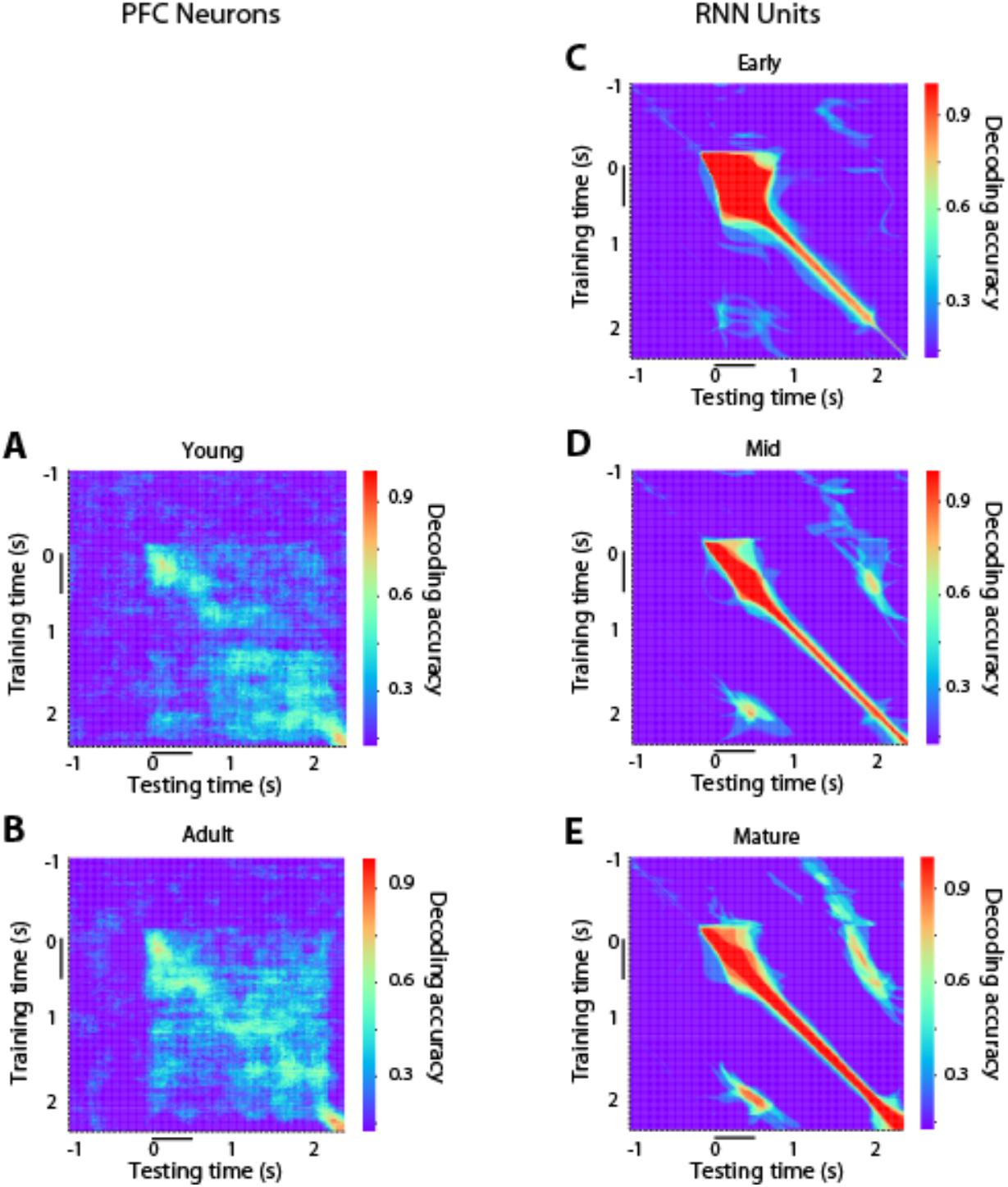
Decoding Accuracy. Cross-temporal decoding accuracy in the ODR task is plotted for PFC data (A-B) and RNN data (C-E). Color plot at each bin represents the accuracy of an SVM decoder trained with a 200 ms bin of data obtained at the time point plotted in the ordinate and tested with the data at the time point indicated in the abscissa. Horizontal bar next to the time scale represents time of cue appearance.

**Supplementary Figure 6.**
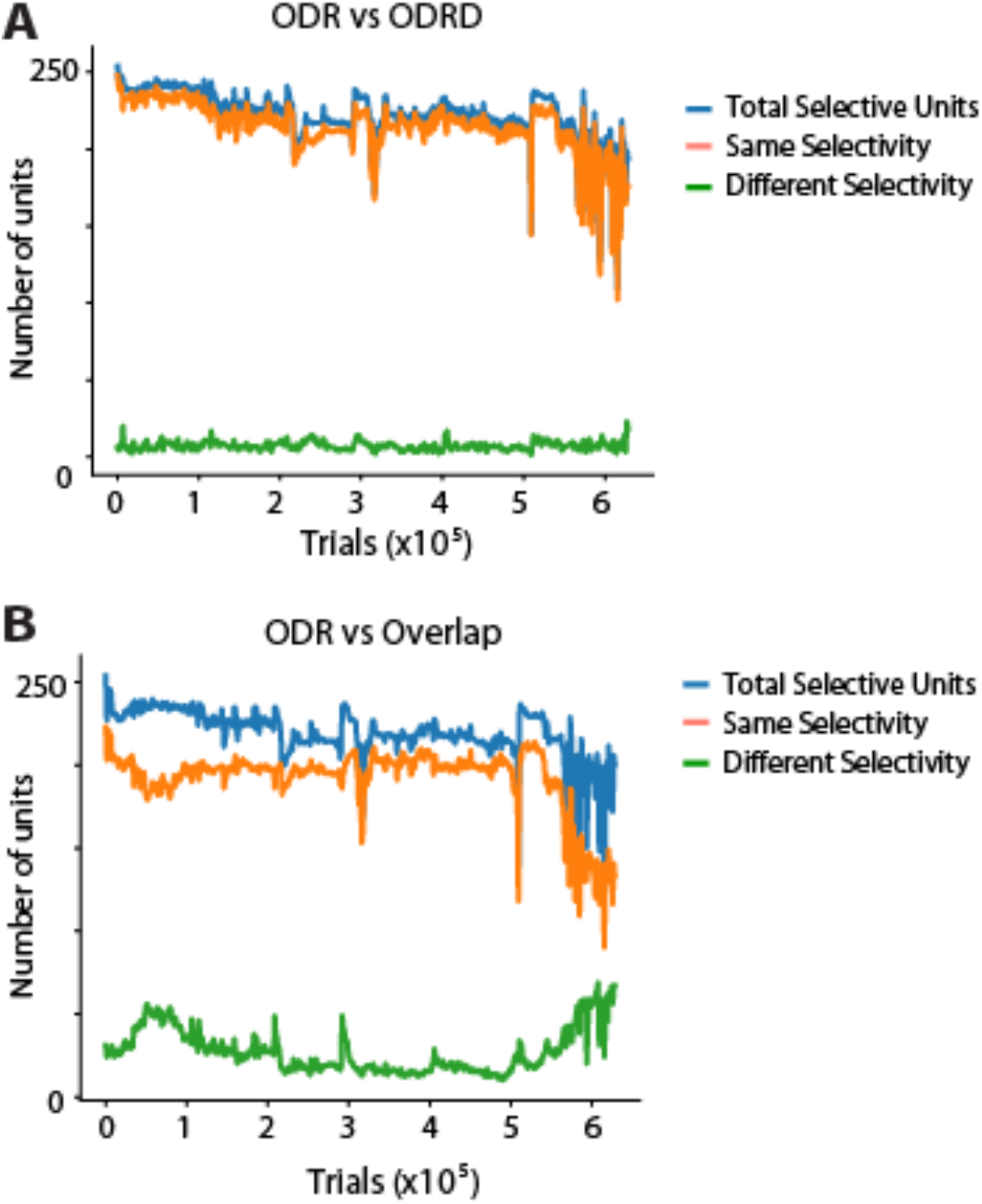
Generalization of RNN activity across tasks. A. Blue line represents number of RNN units (out of 256) exhibiting selectivity for the location of the stimulus during the cue period in the ODR task as a function of trials. Yellow line represents the number of units that exhibit the highest response for the same stimulus location in the ODR task as they due in the ODRD task. Green line represents the number of units that exhibit their best response for different cue locations in the ODR and ODRD tasks. B. Same analysis for the behavior of units in the ODR task, and the Overlap variant of the antisaccade task.

**Supplementary Figure 7.**
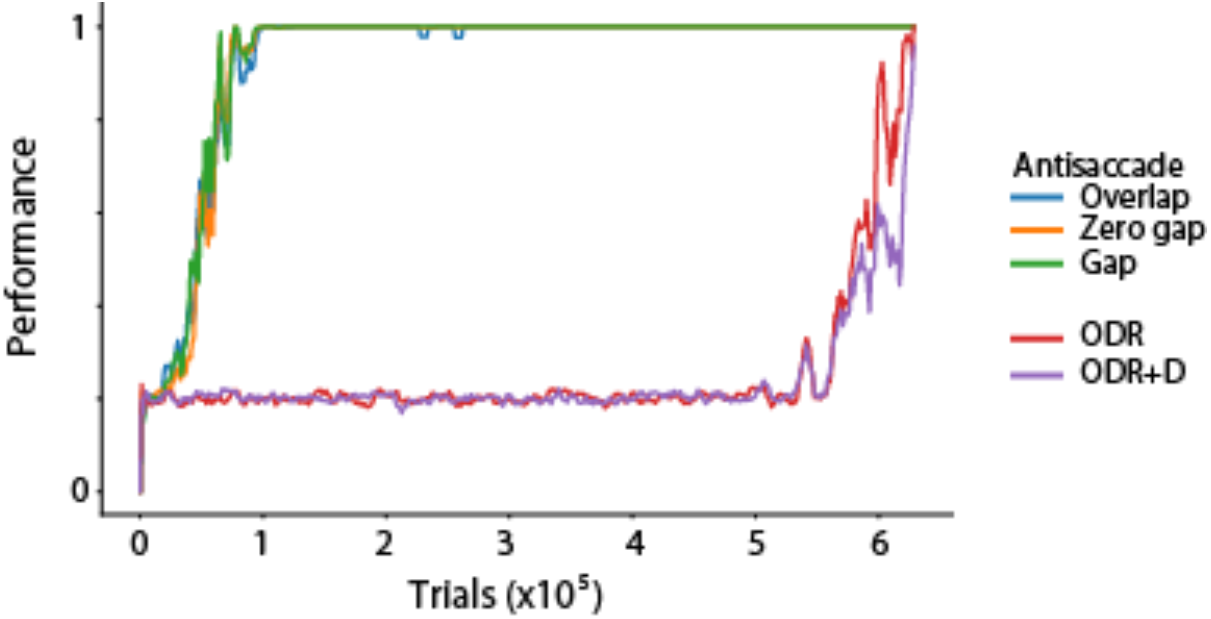
Task performance during training. Performance in the three antisaccade and two ODR tasks are shown as a function of trials, while the network is trained to perform all of them simultaneously.

**Supplementary Figure 8.**
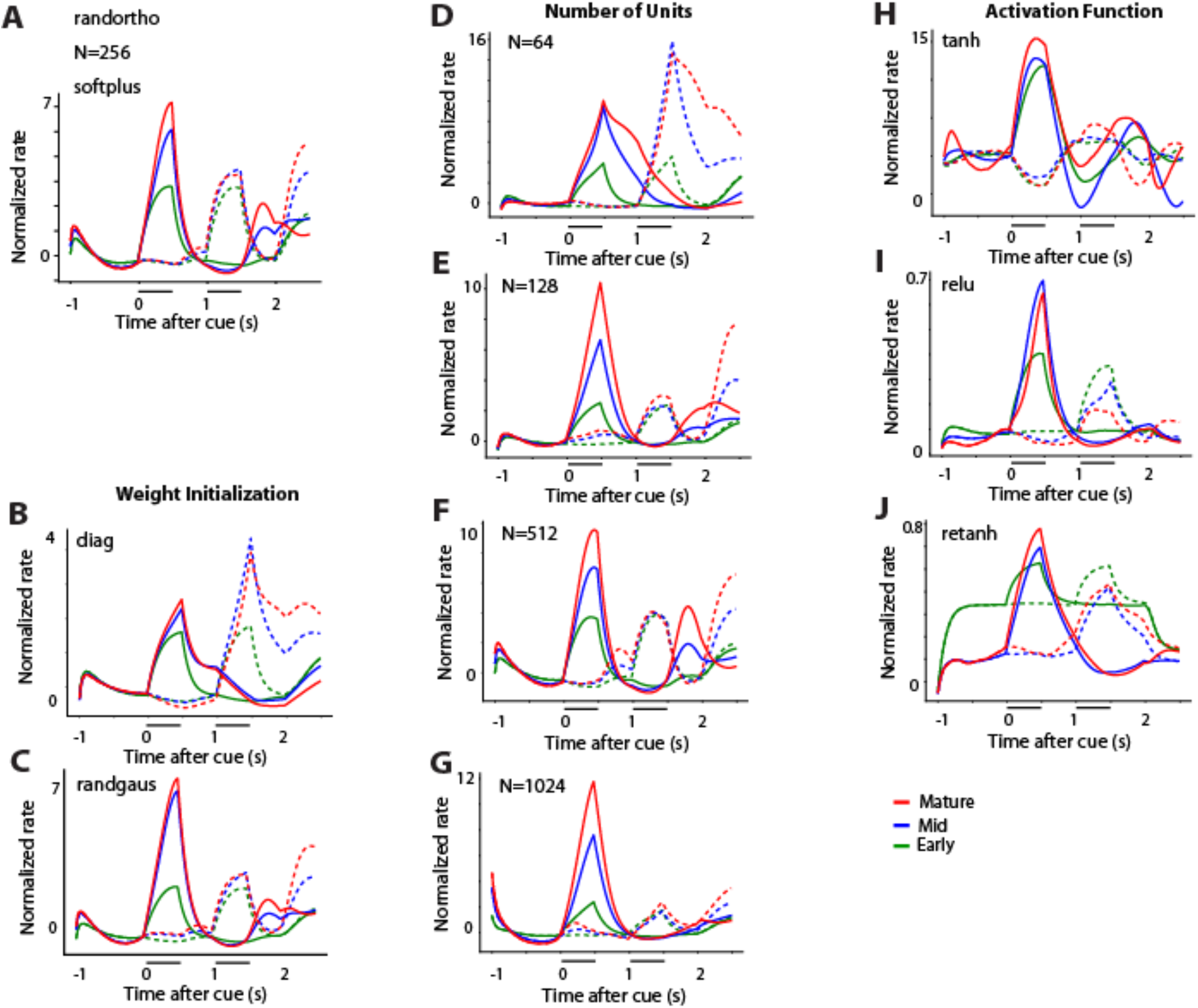
Effect of RNN network hyper-parameters. **A.** Mean firing rate of RNN units in the ODRD task, in the early, mid, and mature networks (same plots as in Fig. 4E-G, superimposed on each other). These simulations were obtained with the parameters indicated in the plot: random orthogonalized initialization, 256 units, and softplus activation function. **B-C**. Networks obtained with different weight initializations: diagonalized (B) and random Gaussian (C). **D-G**. Networks with different numbers of units, varying from 64 to 1024. **H-K**. Networks obtained with different activation functions: hyperbolic tangent (H), rectified linear unit (I), rectified hyperbolic tangent (J).

## Notes

https://github.com/maiziezhoulab/RNN_BrainMaturation

### Competing Interest Statement

The authors have declared no competing interest.

